# Spatial modulation of visual signals arises in cortex with active navigation

**DOI:** 10.1101/832915

**Authors:** E. Mika Diamanti, Charu Bai Reddy, Sylvia Schröder, Tomaso Muzzu, Kenneth D. Harris, Aman B. Saleem, Matteo Carandini

## Abstract

During navigation, the visual responses of neurons in primary visual cortex (V1) are modulated by the animal’s spatial position. Here we show that this spatial modulation is similarly present across multiple higher visual areas but largely absent in the main thalamic pathway into V1. Similar to hippocampus, spatial modulation in visual cortex strengthens with experience and requires engagement in active behavior. Active navigation in a familiar environment, therefore, determines spatial modulation of visual signals starting in the cortex.

## Introduction

There is increasing evidence that the activity of the mouse primary visual cortex (V1) is influenced by navigational signals (Fiser et al., 2016; Fournier et al., 2020; Haggerty and Ji, 2015; Ji and Wilson, 2007; Saleem et al., 2018). During navigation, indeed, the responses of V1 neurons are modulated by the animal’s estimate of spatial position (Saleem et al., 2018). The underlying spatial signals covary with those in hippocampus, and are affected similarly by idiothetic cues (Fournier et al., 2020; Saleem et al., 2018).

It is not known, however, how this spatial modulation varies along the visual pathway. Spatial signals might enter the visual pathway upstream of V1, in the lateral geniculate nucleus (LGN). Indeed, spatial signals have been seen elsewhere in thalamus (Jankowski et al., 2015; Taube, 1995) and possibly also in LGN itself (Hok et al., 2018). Spatial signals might also become stronger downstream of V1, in higher visual areas. For instance, they might be stronger in parietal areas such as A and AM (Hovde et al., 2018), because many neurons in parietal cortex are associated with spatial coding (Krumin et al., 2018; McNaughton et al., 1994; Nitz, 2006; Save and Poucet, 2009; Whitlock et al., 2012; Wilber et al., 2014).

Moreover, it is not known if spatial modulation of visual signals requires experience in the environment and active navigation. In the navigational system, spatial encoding is stronger in active navigation than during passive viewing, when most hippocampal place cells lose their place fields (Chen et al., 2013; Song et al., 2005; Terrazas et al., 2005). In addition, both hippocampal place fields and entorhinal grid patterns grow stronger when an environment becomes familiar (Barry et al., 2012; Frank et al., 2004; Karlsson and Frank, 2008). If spatial modulation signals reach visual cortex from the navigational system, therefore, they should grow with active navigation and with experience of the environment.

## Results

To characterize the influence of spatial position on visual responses, we used a virtual reality (VR) corridor with two visually matching segments (Saleem et al., 2018) (Fig. 1). We used two-photon microscopy to record calcium activity across the visual pathway. Mice were head-fixed and ran on a treadmill to traverse a 1-dimensional virtual corridor made of two visually-matching 40 cm segments each containing the same two visual textures (Fig. 1b *top*). A purely visual neuron would respond similarly in both segments, while a neuron modulated by spatial position could favor one segment.

**Fig. 1.**
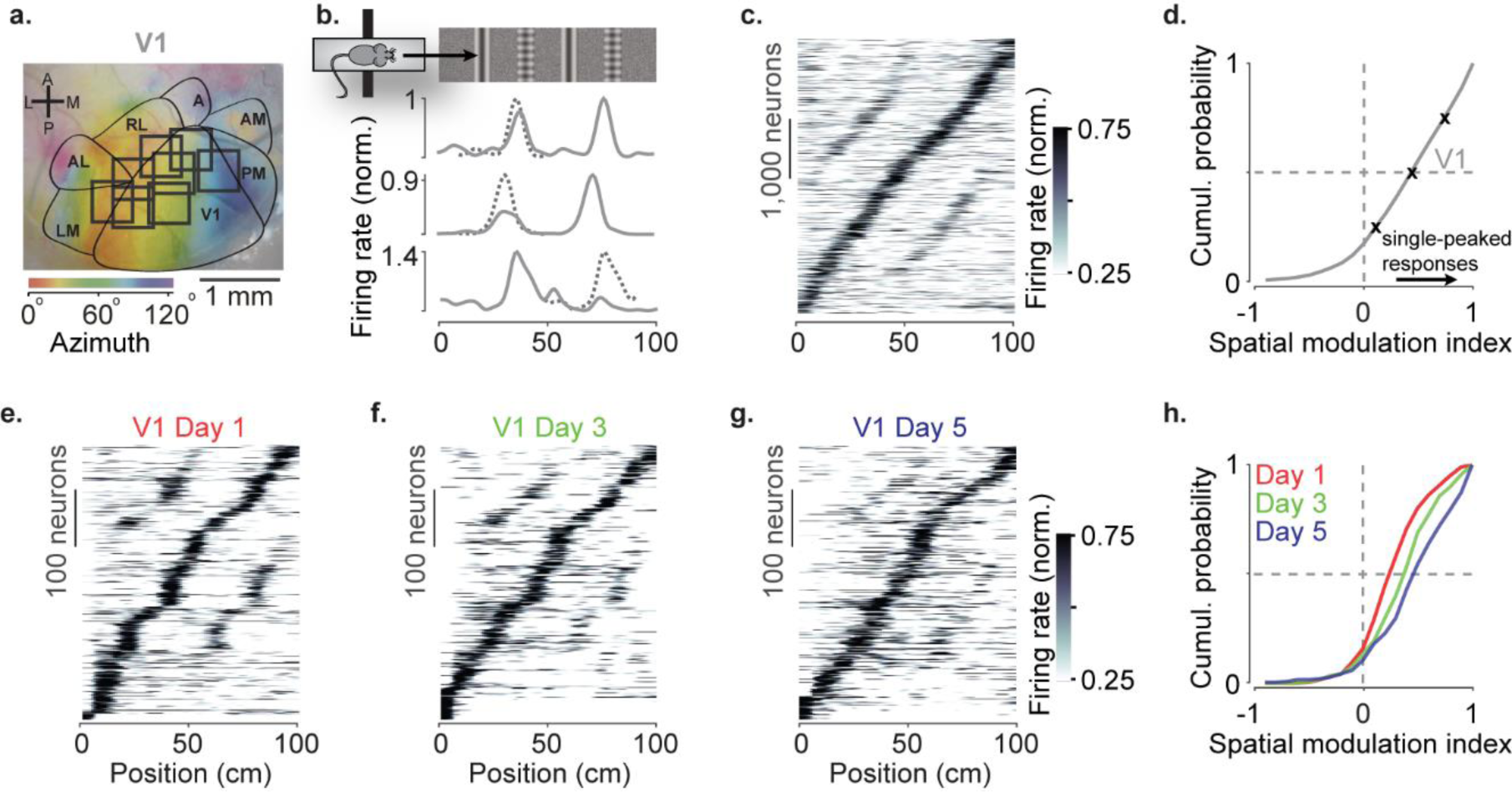
Spatial modulation strengthens with experience. **a**. Example retinotopic map (*colors*) showing borders between visual areas (*contours*) and imaging sessions targeting V1 fully or partly (*squares*, field of view: 500×500 µm). **b**. Normalized firing rate of three example V1 neurons, as a function of position in the virtual corridor. The corridor had two landmarks that repeated after 40 cm, creating visually matching segments (*top*). Dotted lines are predictions assuming identical responses in the two segments. **c**. Firing rates of 4,602 V1 neurons (out of 16,238) whose activity was modulated along the corridor (≥ 5% explained variance), ordered by the position of their peak response. The ordering was based on separate data (odd-numbered trials). **d**. Cumulative distribution of the spatial modulation index (SMI) for the V1 neurons. Only neurons responding within the visually-matching segments are included (2,992 / 4,602). *Crosses* mark the 25^th^, 50^th^, and 75^th^ percentiles and indicate the 3 example cells in **b. e-g** Response profiles obtained from the same field of view in V1 across the first days of experience of the virtual corridor (days 1, 3 and 5 are shown) in two mice. **h**. Cumulative distribution of SMI for those 3 days, showing median SMI growing from 0.24 to 0.38 to 0.45 across days.

As we previously reported, spatial position powerfully modulated the visual responses of V1 neurons (Fig 1a-d). V1 neurons tended to respond more strongly at a single location (Fournier et al., 2020; Saleem et al., 2018) (Fig. 1b) and their preferred locations were broadly distributed along the corridor (Fig. 1c, Supplementary Figure 1a-c). To quantify this spatial modulation of visual responses, we defined a spatial modulation index (SMI) as the normalized difference of responses at the two visually matching positions (preferred minus non–preferred, divided by their sum, with preferred position defined on held-out data). The distribution of SMIs across V1 neurons heavily skewed towards positive values (Fig. 1d), which correspond to a single peak as a function of spatial position. The median SMI for responsive V1 neurons was 0.39 ± 0.19 (n = 39 sessions) and 44% of V1 neurons (1,322/2,992) had SMI>0.5.

Spatial modulation in V1 grew with experience (Fig. 1e-h). In two mice, we measured spatial modulation across the first 5 days of exposure to the virtual corridor, imaging the same V1 field of view across days. Response profiles on Day 1 showed two pronounced peaks (Fig. 1e). By Day 5, response profiles were more single-peaked, and resembled those recorded in experienced mice (Fig. 1c, f, g). Indeed, the spatial modulation increased with experience and was significantly larger on Day 5 compared to Day 1 (Fig. 1h, median SMI: 0.45 on Day 5 vs. 0.24 on Day 1; p < 10^−12^, two-sided Wilcoxon rank sum test).

In contrast to V1 neurons, spatial position barely affected the visual responses of LGN afferents in experienced mice (Fig. 2a-d). LGN boutons in layer 4 gave similar visual responses in the two segments of the corridor (Fig. 2b,c) and the locations where they fired clustered around the landmarks as expected from purely visual responses (Supplementary Figure 1e). The spatial modulation in LGN boutons was small (Fig. 2d). The median SMI across sessions for responsive LGN boutons was barely above zero (0.07 ± 0.05, n = 19 sessions), markedly smaller than the SMIs of V1 neurons (p = 10^−5^, left-tailed two-sample *t*-test). Only 4% of LGN boutons (37/842) had SMI > 0.5, compared to 44% in V1. Moreover, the preferred position of those few neurons often fell in the first half of the corridor as would be expected from contrast adaptation mechanisms (Dhruv and Carandini, 2014) (Supplementary Figure 1f).

**Fig. 2.**
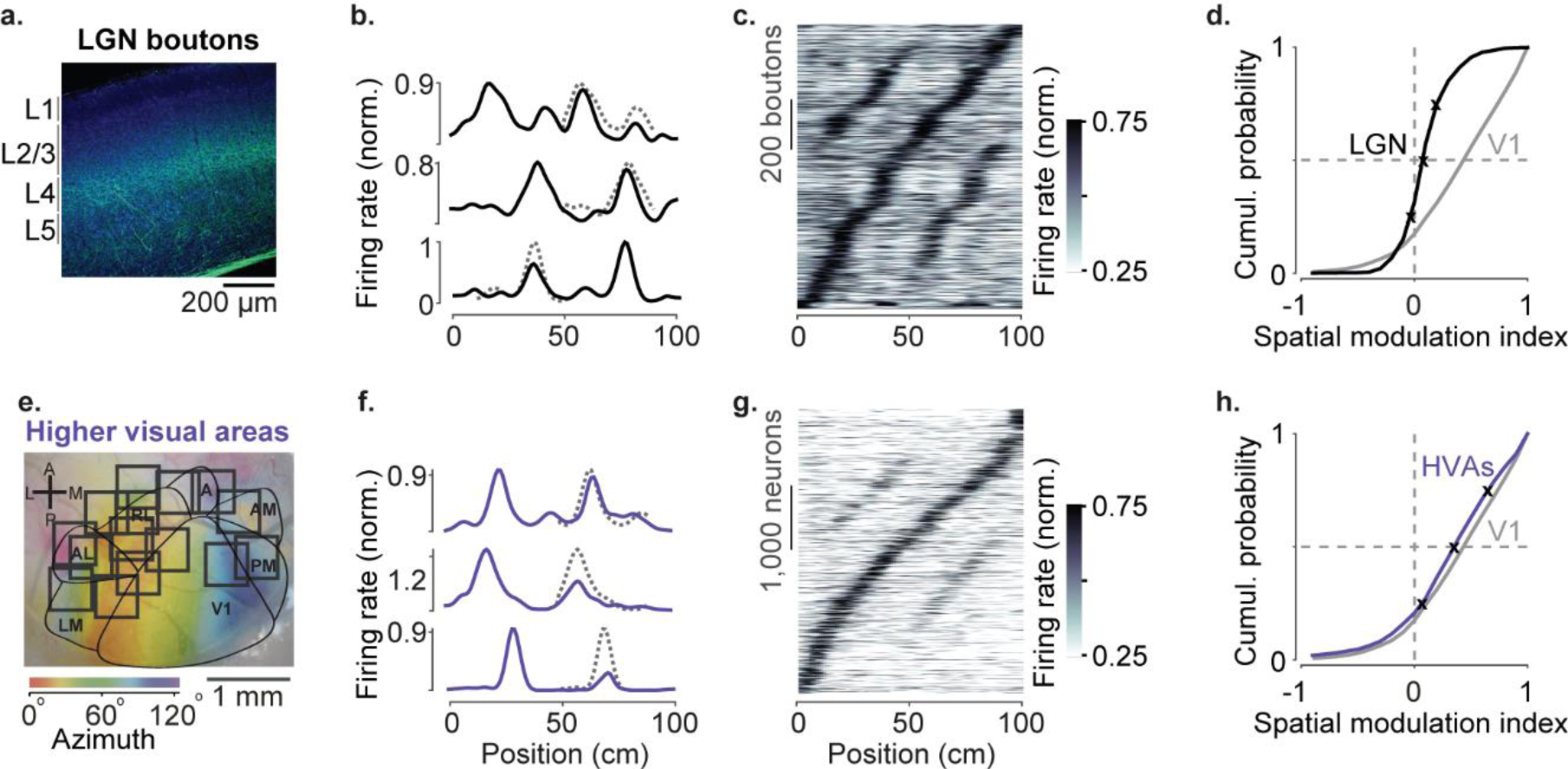
Modulation of visual responses along the visual pathway during navigation. **a**. Confocal images of LGN boutons expressing GCaMP (GFP; *green*) among V1 neurons (Nissl stain; *blue*). GCaMP expression is densest in layer 4 (L4). **b**. Normalized firing rate of three example LGN boutons, as a function of position in the virtual corridor. Dotted lines are predictions assuming identical responses in the two segments. **c**. Firing rates of 1,140 LGN boutons (out of 3,182) whose activity was modulated along the corridor (≥ 5% explained variance), ordered by the position of their peak response. The ordering was based on separate data (odd-numbered trials). **d**. Cumulative distribution of the spatial modulation index (SMI) for the LGN boutons. Only boutons responding within the visually-matching segments are included (LGN: 840/1,140). *Crosses* mark the 25^th^, 50^th^, and 75^th^ percentiles and indicate the 3 example cells in **b. e**. Same as in Fig. 1a, showing imaging sessions targeting fully or partly 6 higher visual areas (HVA). **f**-**h**. Same as **b**-**d**, showing response profiles of HVA neurons (**g:** 4,381 of 18,142 HVA neurons; **h**: 2,453 of those neurons).

Similar results were seen in recordings from LGN neurons (Supplementary Figure 2). We performed extracellular electrophysiology recordings in LGN (2 mice, 5 sessions). LGN units gave responses and SMI similar to LGN boutons (0.06 ± 0.02, p = 0.92, two-sample *t*-test) and thus markedly different from V1 (p = 0.004, left-tailed two-sample *t*-test).

Spatial modulation was broadly similar across higher visual areas, and not significantly larger than in V1 (Fig. 2e-h, Fig. 3). We measured activity in six visual areas that surround V1 (LM, AL, RL, A, AM, and PM), and found strong modulation by spatial position (Fig. 2f,g). Pooling across these areas, the median SMI across sessions was 0.40 ± 0.12, significantly larger than zero (n = 52 sessions, p = 10^−10^, right-tailed Wilcoxon signed rank test, Fig. 2h) and not significantly different from V1 (two-sample *t*-test: p = 0.83). Spatial modulation was present in each of the six areas (Fig. 3), and as in V1 (Saleem et al., 2018) it could not be explained by other factors such as running speed, reward events, pupil size and eye position (Supplementary Figure 3).

**Fig. 3.**
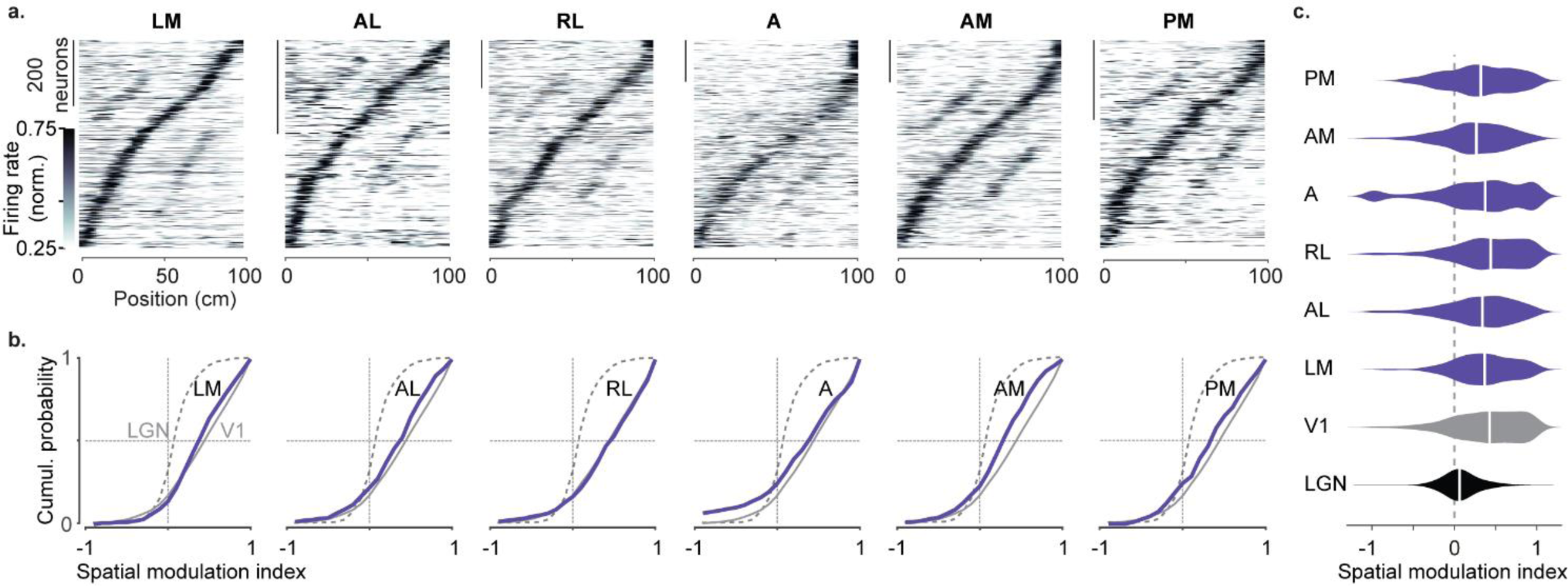
Spatial modulation of individual higher visual areas. **a**. Response profile patterns obtained from even trials (ordered and normalized based on odd trials) for 6 visual areas. Only response profiles with variance explained ≥ 5% are included (LM: 629/1503 AL: 443/1774 RL: 866/5192 A: 997/4126 AM: 982/3278 PM: 519/2509). **b**. Cumulative distribution of the spatial modulation index in even trials for each higher visual area (purple). Dotted line: LGN (same as in Fig. 2), Gray: V1 (same as in Fig. 1). **c**. Violin plots showing the SMI distribution and median SMI (white vertical line) for each visual area (median ± m.a.d. LGN: 0.07±0.11; V1: 0.43±0.31; LM: 0.37±0.25; AL: 0.34±0.28; RL: 0.44±0.31; A: 0.37±0.34; AM: 0.27±0.26; PM: 0.32±0.32).

We observed small differences in spatial modulation between areas, which may arise from biases in retinotopy combined with the layout of the visual scene. Visual patterns in the central visual field were further away in the corridor, and thus were likely less effective in driving responses than patterns in the periphery, which were closer to the animal and thus larger. In V1, spatial modulation was larger in neurons with central rather than peripheral receptive fields (Supplementary Figure 4). A similar trend was seen across higher areas, with slightly lower SMI in areas biased towards the periphery (AM, PM) (Garrett et al., 2014; Wang and Burkhalter, 2007; Zhuang et al., 2017) than in areas biased towards the central visual field (LM, RL, Fig. 3c).

We next asked whether visual responses would be similarly modulated when animals passively viewed the environment. After recordings in virtual reality (‘VR’), we played back the same video regardless of the mouse’s movements (‘replay’). We separated data taken during running (running speed > 1 cm/s in at least 10 trials, ‘running replay’), and rest (‘stationary replay’) periods.

Passive viewing affected the baseline activity of LGN boutons but not their spatial modulation, which remained negligible in all conditions (Fig. 4a-d). During ‘running replay’ the baseline activity of LGN boutons decreased slightly (Fig. 4a,b, p = 0.003, paired-sample right-tailed t-test, Supplementary Figure 5a). However, their mean SMI in ‘running replay’ remained a mere 0.06 ± 0.02 (s.e., n = 18 sessions), not significantly different from the 0.09 ± 0.03 measured in VR (Fig. 4b, p = 0.12, paired-sample right-tailed *t*-test). Similar results were obtained during stationary replay (Fig. 4c-d): baseline activity decreased markedly (Erisken et al., 2014) (Fig. 4d,e, p = 10^−65^ paired-sample right-tailed t-test, Supplementary Figure 5b), but the SMI remained negligible at 0.03 ± 0.01 and not larger than the 0.07 ± 0.02 measured in VR (n = 18 sessions, p = 0.05, Fig. 4d).

**Fig. 4.**
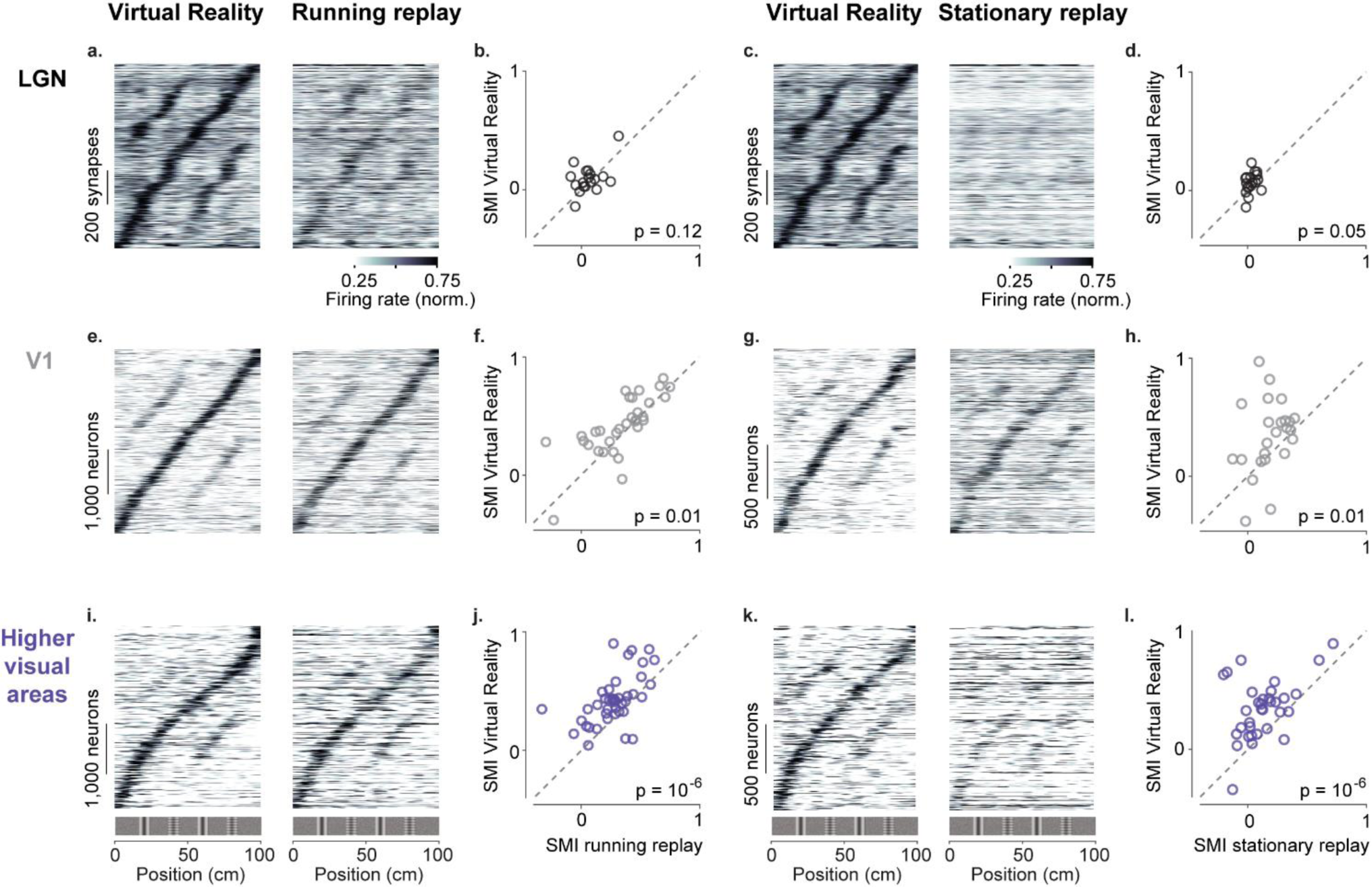
Active navigation enhances modulation by spatial position in visual cortical areas. **a**. Response profiles of LGN boutons in Virtual Reality (*left*) that also met the conditions for running replay (*right*; at least 10 running trials per recording session), estimated as in Fig. 1c. Response profiles of LGN boutons during running replay were ordered by the position of their maximum response estimated from odd trials in Virtual Reality (same order and normalization as in *left* panel). **b**. Median spatial modulation index (SMI) per recording session in Virtual Reality versus running replay for LGN (each circle corresponds to a single session; p values from paired-sample right-tailed *t*-test). **c, d**. Same as **a**,**b** for stationary replay. **e-h**. Same as in **a-d** for V1 neurons. **i-l**. Same as in **a-d** for neurons in higher visual areas.

Many V1 neurons showed weaker modulation by spatial position during replay than in VR, particularly during stationary replay (Fig. 4e-h). In V1, ‘running replay’ reduced mean SMI by ∼20%, from 0.42 ± 0.04 in VR to 0.34 ± 0.04 (s.e., n = 32 sessions, p = 0.01, Fig. 4e-f). This decrease in spatial modulation was not associated with firing rate differences (p > 0.05, Supplementary Figure 5a). Therefore, running without matched visual feedback did not result in the same spatial modulation as active navigation. The decrease in spatial modulation was even greater during rest (‘stationary replay’, Fig. 4g-h). As expected in mice that are not running, firing rates decreased markedly (Keller et al., 2012; Saleem et al., 2013)(p = 10^−24^, Supplementary Figure 5b). In addition, the mean SMI almost halved from 0.34 ± 0.06 in VR to 0.18 ± 0.03 in stationary replay, a significant decrease (p = 0.01, n = 24 sessions).

These effects were especially marked in higher visual areas (Fig. 4i-l). Here, ‘running replay’ reduced mean SMI by ∼37%, from 0.41 ± 0.03 in VR to 0.26 ± 0.03 (s.e., n = 41 sessions, p = 10^−6^, Fig. 4i,j), without affecting firing rates (p > 0.05 in all areas, Supplementary Figure 5a). Even stronger effects were seen during stationary replay (Fig. 4k,l), where the mean SMI decreased ∼65%, from 0.34 ± 0.04 in VR to 0.12 ± 0.04 (p = 10^−6^, n = 33 sessions, Fig. 4k,l). This effect was accompanied by decreased firing in some areas, notably AM (p = 0.03) and PM (p = 10^−09^ Supplementary Figure 5b).

## Discussion

Taken together, these results indicate that upon experience active navigation shapes responses along the visual pathway starting in the cortex.

Navigational signals in V1 strengthened across the first few days, at slower timescales than in hippocampus (Frank et al., 2004; Karlsson and Frank, 2008) but similar to retrosplenial cortex (Mao et al., 2018). Similar results have been observed when decoding spatial position from V1 across days of exposure to a slightly changing environment (Fiser et al., 2016).

Navigational signals in V1 are not inherited from thalamic inputs from LGN, as this modulation was practically absent in LGN inputs to layer 4 and in LGN neurons themselves (regardless of what layer they project to (Cruz-Martín et al., 2014)). However, our mice were head restrained and hence lacked vestibular inputs (Ravassard et al., 2013; Russell et al., 2006). Perhaps when mice freely move, LGN does show some spatial modulation (Hok et al., 2018), which is possibly amplified in V1 by non-linear mechanisms (Chariker et al., 2016; Lien and Scanziani, 2013).

Navigational signals affected all cortical visual areas approximately equally, consistent with the widespread neural coding of task-related information across the posterior cortex (Koay et al., 2019; Minderer et al., 2019). In addition, all areas gave stronger responses during active behavior than during passive viewing.

Navigational signals may reach visual cortex through retrosplenial cortex (Makino and Komiyama, 2015), an area that contains experience-dependent spatial signals (Mao et al., 2017, 2018) and is more strongly modulated by active navigation than V1 (Fischer et al., 2020). Another candidate is anterior cingulate cortex (Zhang et al., 2014), whose dense projections to V1 carry signals related to locomotion (Leinweber et al., 2017). The route that navigational signals take across the cortex is yet to be charted.

## Acknowledgements

We thank Julien Fournier for helpful discussions, Michael Krumin for assistance with two-photon imaging, and Karolina Socha for advice on imaging LGN boutons. This work was supported by EPSRC (PhD award F500351/1351 to E.M.D.), by a Wellcome Trust/Royal Society Sir Henry Dale Fellowship (200501 to A.B.S.), by a Human Frontiers in Science Program (grant RGY0076/2018 to A.B.S.), and by the Wellcome Trust (grant 205093 to M.C. and K.D.H.). M.C. holds the GlaxoSmithKline / Fight for Sight Chair in Visual Neuroscience.

## Author contributions

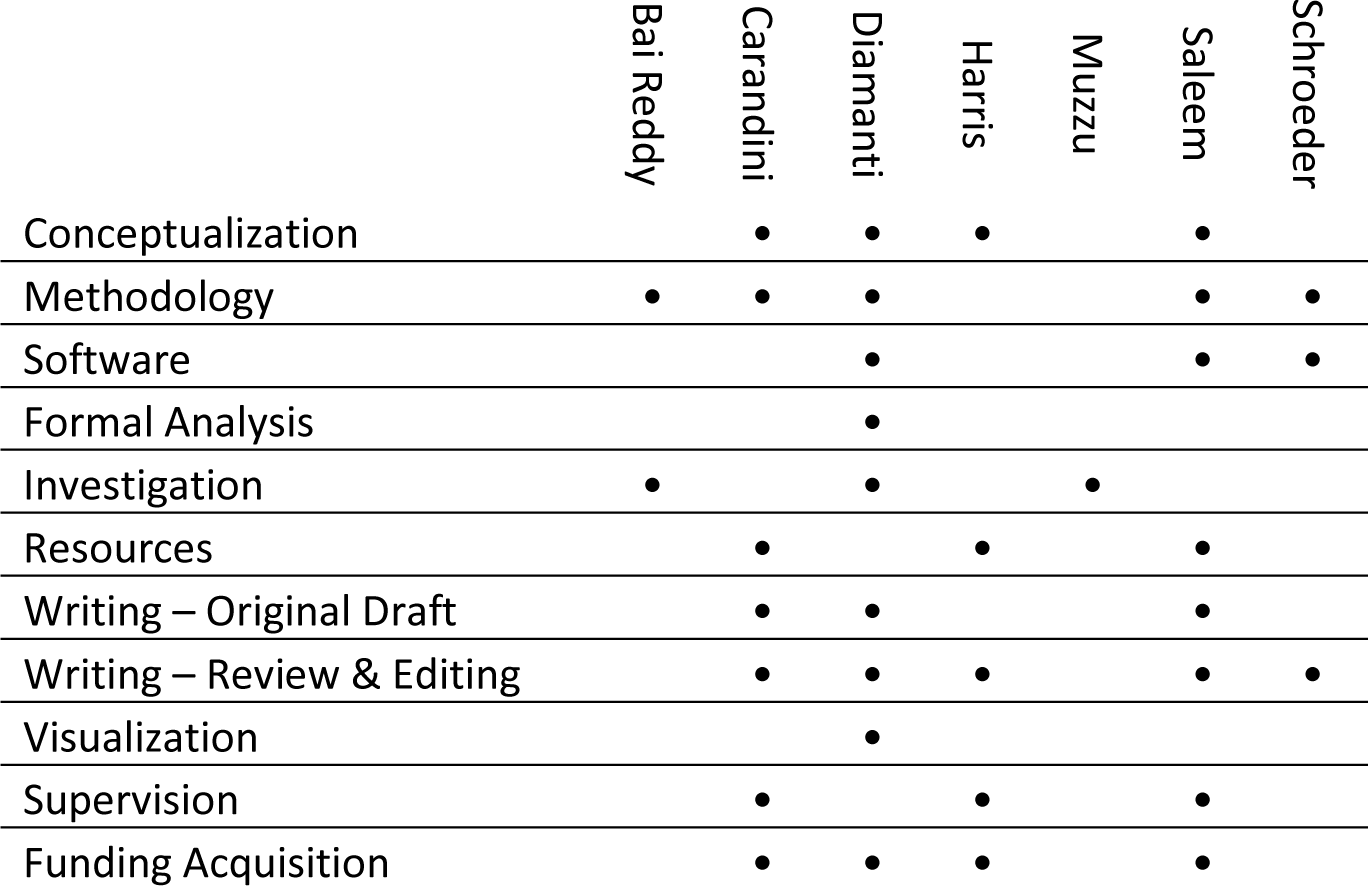

## Supplementary Figures

**Supplementary Figure 1.**
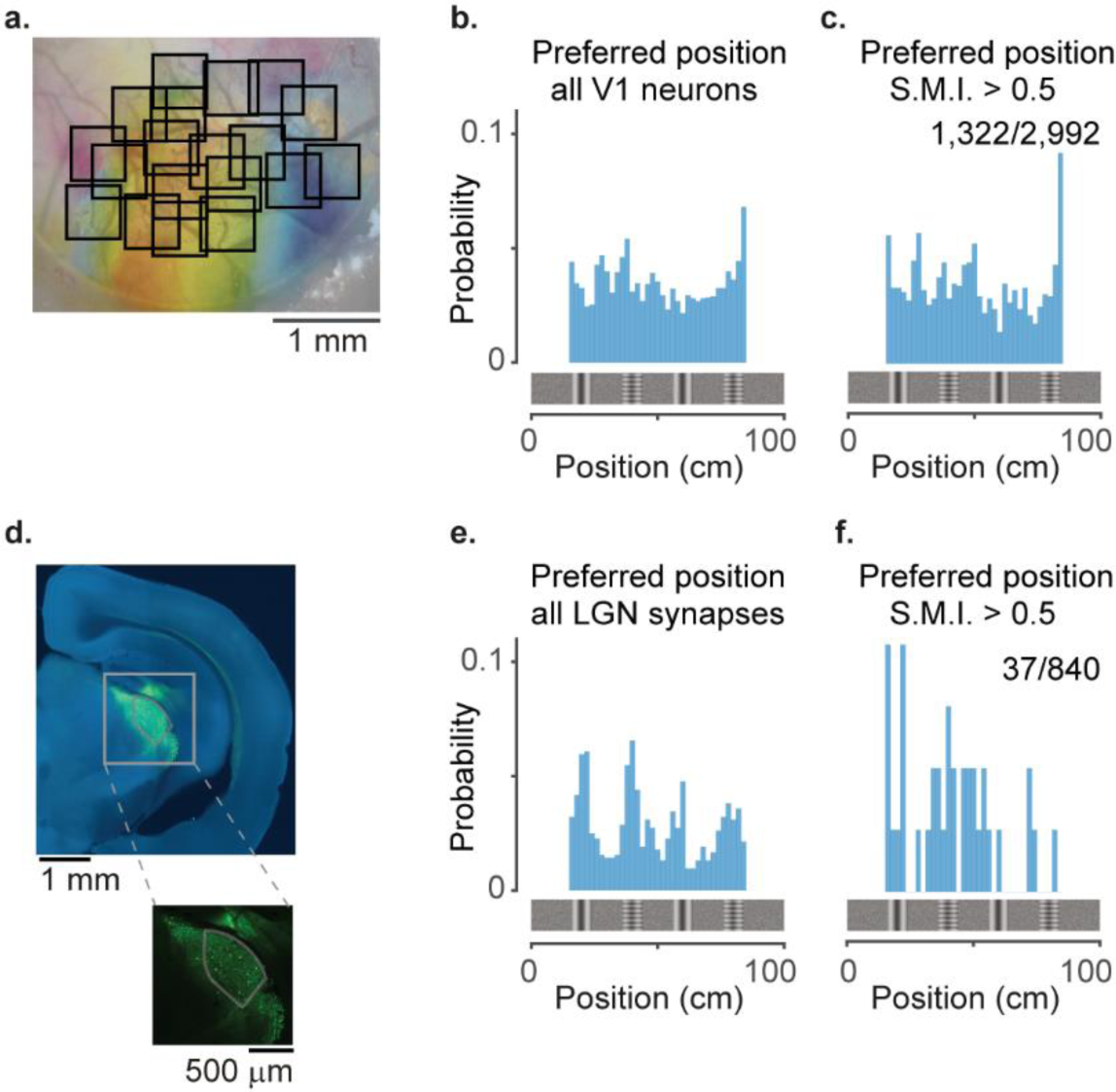
LGN boutons tile up the virtual corridor differently from V1 neurons. **a**. Two-photon imaging of cell bodies in visual cortex across multiple days: example retinotopic map with single-session fields of view (*black squares*). Combination of Fig. 1a and Fig. 2e. **b**. Distribution along the corridor of preferred positions within the visually matching segments for all V1 neurons (bin size: 1 cm) **c**. Same as in **b**. for V1 neurons with S.M.I. > 0.5 (n = 1,322/2,992). **d**. Example image showing viral expression of GCaMP (*green*) in the visual thalamus of the right hemisphere (stained with Nissl in *blue*). *Inset:* zoomed-in image showing GCaMP expression in LGN and the surrounding nuclei. The LGN border (*gray contour*) was determined using SHARP-Track (Shamash et al., 2018). **e**. Distribution of preferred positions along the corridor within the visually matching segments for all 840 LGN boutons (bin size: 1 cm). **f**. Same as in **e**. for LGN boutons with SMI > 0.5 (n = 37/840).

**Supplementary Figure 2.**
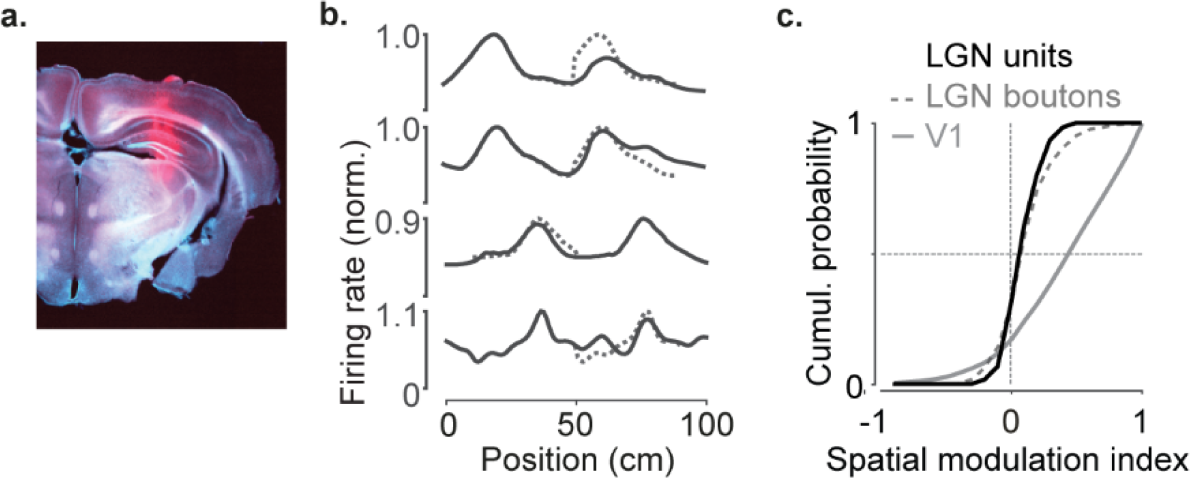
LGN boutons and units give similar responses. **a**. Example reconstructed electrode tracks terminating in LGN (red: DiI; brain image stained with *DAPI*). **b**. Response profile patterns of example LGN units recorded using extracellular multi-electrode arrays. Responses are from even trials, ordered and normalized from odd trials. Dotted lines are predictions assuming identical responses in the two segments. **c**. Cumulative distribution of the spatial modulation index in even trials for LGN units (*black*), LGN boutons (*dotted gray line*, similar to Fig. 2d) and V1 (*gray line*, similar to Fig. 1d) (79 units from 2 animals).

**Supplementary Figure 3.**
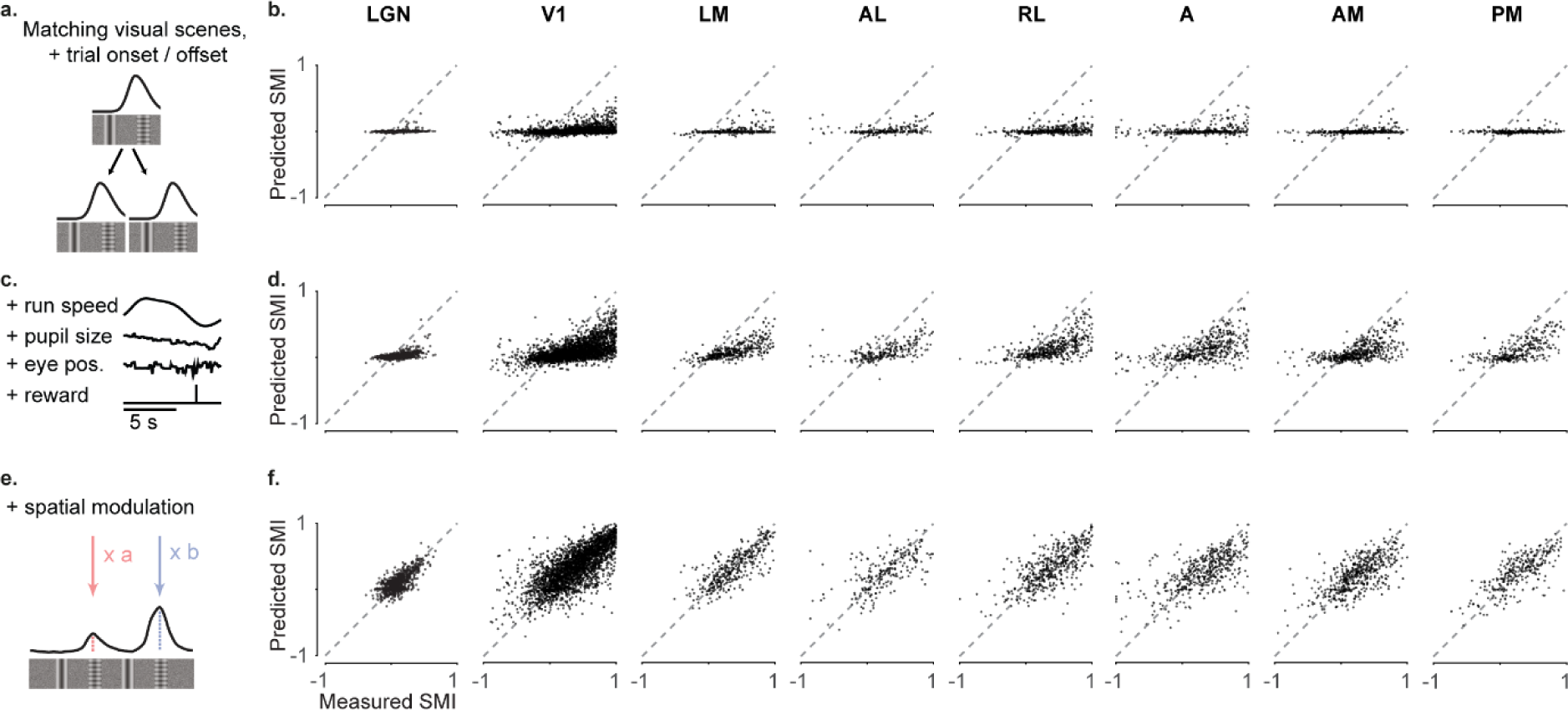
Spatial modulation is not explained by other behavioral and visual factors. We constructed three models to predict the activity of individual neurons from successively larger sets of predictor variables(Saleem et al., 2018) **a, b**. Measured spatial modulation index for each visual area versus predictions of the simplest model (‘purely visual model’). The ‘purely visual model’ considers only the repetitions of the visual scenes, trial onset and offset, and as expected, fails to predict the SMIs estimated from the data. Each point represents a neuron. **c, d**. Measured spatial modulation index for each cortical visual area versus predictions of the ‘non-spatial’ model. The ‘non-spatial’ model also includes the contribution by behavioral factors that can differ within and across trials: speed, reward times, pupil size, and eye position. Adding these factors improves predictions compared to the ‘purely visual’ model but fails to match the measured SMIs. Therefore, the joint contribution of all task-related and visual factors is not sufficient to explain the observed spatial modulation **e, f**. Measured spatial modulation index for each visual area versus predictions of the ‘spatial’ model. The spatial model allows the peaks in the visually matching segments to vary independently. It provides a much better match to the data.

**Supplementary Figure 4.**
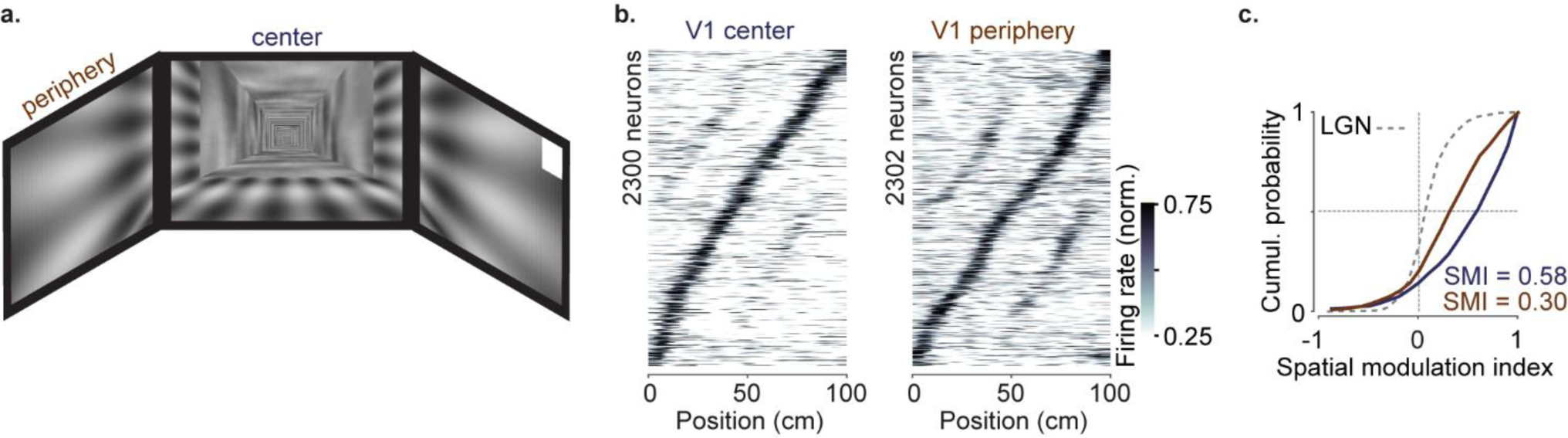
Neurons with central receptive fields showed stronger spatial modulation than neurons with peripheral receptive fields due to the layout of the visual scenes. a. Cartoon of the virtual reality scenes layout. b. Response profile patterns obtained from even trials (ordered and normalized based on odd trials) for portions of V1 with average receptive fields in the center (‘V1 center’; <40 deg azimuth angle; *left*) or in the periphery (‘V1 periphery’; *right*). c. Cumulative distribution of the spatial modulation index in even trials for ‘V1 center’ (*purple*) or ‘V1 periphery’ (*orange*); *dotted line:* Distribution of LGN boutons (same as in Fig. 2d).

**Supplementary Figure 5.**
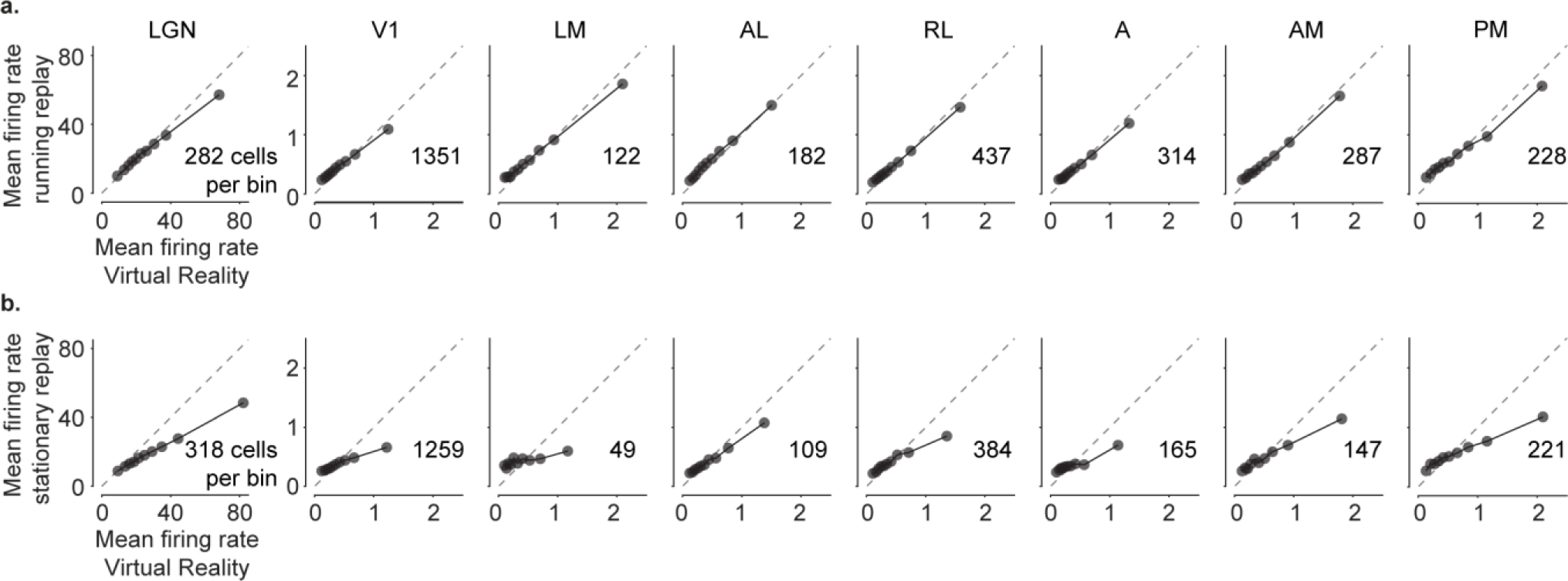
Comparison of firing rates in Virtual Reality and replay conditions. **a**. Comparison of the mean firing rate between Virtual Reality and running replay for all probed visual areas, split into 10 percentile bins. **b**. Same as in **a**. for stationary replay. Mean firing rates were similar between Virtual Reality and running replay but decreased significantly during stationary replay in many areas. (paired-sample right-tailed *t*-test: running replay: LGN: p = 0.003; p > 0.05 in all cortical areas; stationary replay: LGN: 10^−65^, V1: 10^−24^, LM: 0.93, AL: 0.15, RL: 0.06, A: 0.56, AM: 0.03, PM: 10^−09^).

## Methods

All experimental procedures were conducted under personal and project licenses issued by the Home Office, in accordance with the UK Animals (Scientific Procedures) Act 1986.

For calcium imaging experiments in visual cortex, we used double or triple transgenic mice expressing GCaMP6 in excitatory neurons (5 females, 1 male, implanted at 4-10 weeks). The triple transgenics expressed GCaMP6 fast (Madisen et al., 2015) (Emx1-Cre;Camk2a-tTA;Ai93, 3 mice). The double transgenic expressed GCaMP6 slow (Wekselblatt et al., 2016) (Camk2a-tTA;tetO-G6s, 1 mouse). Because Ai93 mice may exhibit aberrant cortical activity (Steinmetz et al., 2017), we used the GCamp6 slow mice to validate the results obtained from the GCaMP6 fast mice. For calcium imaging experiments of LGN boutons, we used 3 C57BL/6 mice (3 females, implanted at 6-9 weeks).

### Surgical procedures

For two-photon calcium imaging of activity in visual cortex, 4–10–week–old mice were implanted with an 8 mm circular chamber and a 4 mm craniotomy was performed over the left or right visual cortex as previously described (Saleem et al., 2018). The craniotomy was performed by repeatedly rotating a biopsy punch and it was shielded with a double coverslip (4 mm inner diameter; 5 mm outer diameter).

For two-photon calcium imaging of activity of LGN boutons, 4–10–week–old mice were first implanted with an 8 mm head plate and a 4mm craniotomy was performed over the right hemisphere, as described above. We next injected 253 nL (2.3 nL pulses with an inter-pulse interval of 5 s, total of 110 pulses) of virus AAV9.CamkII.GCamp6f.WPRE.SV40 (5.0×10^12^ GC/ml) into the visual thalamus. To target LGN in the right hemisphere, the pipette was directed at 2.6 mm below the brain surface, 2.3 mm posterior and 2.25 mm lateral from bregma. To prevent backflow, the pipette was kept in place for 5 min after the end of the injection. In addition to dorsal LGN, the virus could infect neighboring thalamic nuclei, including the higher-order visual thalamic nucleus LP, which projects to layer 1 of visual cortex (Roth et al., 2016). Therefore, we imaged boutons only in layer 4, the main recipient of dorsal LGN inputs.

### Perfusion and Histology

Mice were perfused with 4% Paraformaldehyde (PFA) and the brain was extracted and fixed for 24 hours at 4°C in 4% PFA, then transferred to 30% sucrose in PBS at 4° C. The brain was mounted on a benchtop microtome and sectioned at 60 µm slice thickness. Free-floating sections were washed in PBS, mounted on glass adhesion slides, stained with DAPI (Vector Laboratories, H-1500) and cover with a glass-slip before being imaged.

Brain sections were initially imaged on a Zeiss AxioScan with a 4x/0.2-NA objective (Nikon CFI Plan Apochromat Lambda) by stitching 3×3 fields of view (total image size: 6 mm x 6 mm). Images were obtained in two colors: green for GCaMP and blue for DAPI.

To obtain higher magnification images of GCaMP expression in LGN or of LGN boutons in V1 we performed confocal microscopy on a Zeiss LSM 880 with Airyscan with a 10x/0.3-NA objective (Zeiss EC Plan-Neofluar). We imaged GCaMP using an excitation wavelength of 488 nm and DAPI using 405 nm, and averaged across 16 frames.

### Two-photon imaging

Two-photon imaging was performed with a standard multiphoton imaging system (Bergamo II; Thorlabs Inc.) controlled by ScanImage4 (Pologruto et al., 2003). A 970 nm or 920 nm laser beam, emitted by a Ti:Sapphire Laser (Chameleon Vision, Coherent), was targeted onto L2/3 neurons or L4 LGN boutons through a 16x water-immersion objective (0.8 NA, Nikon). Fluorescence signal was transmitted by a dichroic beam splitter and amplified by photomultiplier tubes (GaAsP, Hamamatsu). The emission light path between the focal plane and the objective was shielded with a custom-made plastic cone, to prevent contamination from the monitors’ light.

Multiple-plane imaging was enabled by a piezo focusing device (P-725.4CA PIFOC, Physik Instrumente), and an electro-optical modulator (M350-80LA, Conoptics Inc.) which allowed adjustment of the laser power with depth.

For experiments monitoring activity in visual cortex, we imaged 4 planes set apart by 40 μm. Images of 512⨯512 pixels, corresponding to a field of view of 500×500 μm, were acquired at a frame rate of 30 Hz (7.5 Hz per plane). For experiments monitoring activity of LGN boutons, we imaged 7-10 planes set apart by 1.8 μm (2 to 3 of these planes were fly-back). Images of 256×256 pixels, corresponding to a field of view of 100×100 μm, were acquired at a frame rate of 58.8 Hz.

For experiments in naïve mice (Fig. 1e-h) we targeted the same field of view based on vasculature and recorded from similar depths. We did not attempt to track the same individual neurons across days.

### Widefield calcium imaging

To obtain retinotopic maps we used wide-field calcium imaging, as previously described (Saleem et al., 2018). Briefly, we used a standard epi-illumination imaging system (Carandini et al., 2015; Ratzlaff and Grinvald, 1991) together with an SCMOS camera (pco.edge, PCO AG). A 14°-wide vertical window containing a vertical grating (spatial frequency 0.15 cycles/°), swept (Kalatsky and Stryker, 2003; Yang et al., 2007) the horizontal position of the wind window over 135° of azimuth angle, at a frequency of 2 Hz. To obtain maps for preferred azimuth we combined responses to the 2 stimuli moving in opposite direction (Kalatsky and Stryker, 2003).

### Neuropil receptive fields

To obtain neuropil receptive fields, on each recording session we presented sparse uncorrelated noise for 5 min. The screen was divided into a grid of squares of 4 x 4 degrees size. Each square was turned on and off randomly at a 10 Hz rate. At each moment in time, 2% of the squares were on. To compute the neuropil receptive fields, the field of view was segmented into 5×5 patches (100 µm x 100 µm surface per patch). For each patch, we first averaged the raw fluorescence across the patch’s pixels. We then computed the stimulus-triggered average of the averaged raw fluorescence trace. The response was further smoothed in space and its peak was defined as the patch’s receptive field center.

### Virtual Reality environment

Animals were head-restrained in the center of three LCD monitors (IIyama ProLite E1980SD 19’’) or three 10-inch LCD screens (LP097QX1-SPAV 9.7’’, XiongYi Technology Co., Ltd.) placed at 90° angle to each other. The distance from each screen was 19 cm for the LCD monitors, or 11cm for the LCD screen, so that visual scenes covered the visual field by 135° in azimuth and 42° in elevation.

The Virtual Reality environment was a corridor with two visually matching segments (Saleem et al., 2018). Briefly, the corridor was 8 cm wide and 100 cm long. A vertical grating or a plaid, 8 cm wide each, alternated in the sequence grating-plaid-grating-plaid at 20, 40, 60 and 80 cm from the start of the corridor.

In Virtual Reality mode, animals traversed the virtual environment by walking on a polystyrene wheel (15 cm wide, 18 cm diameter) which allowed movement along a single dimension (forwards or backwards). Running speed was measured online with a rotary encoder (2400 pulses/rotation, Kübler, Germany) and was used to control the update of visual scenes. Upon reaching the 100^th^ cm of the corridor, animals were presented with a gray screen for an inter-trial period of 3 to 5 s, after which they were teleported back to the beginning of the corridor for the next trial. The duration of each trial depended on how long it took the animal to reach the end of the corridor, typically less than 8 s. Trials in which animals did not reach the end of the corridor within 30 s were timed-out and excluded from further analysis. A typical session consisted of more than 50 trials.

In the replay mode, mice were presented with a previous closed-loop session, while still free to run on the wheel.

### Electrophysiological recordings

Mice were implanted with a custom-built stainless-steel metal plate on the skull under isoflurane anesthesia. The area above the right LGN was kept accessible for electrophysiological recordings. Mice were acclimatized to the VR environment for over 5 days. The virtual reality was projected onto a truncated spherical screen, and the mice run through it by running on a 10cm radius Styrofoam (Schmidt-Hieber and Häusser, 2013), using the custom MATLAB code described above (Saleem et al., 2018). 12-24 hours before the first recording, a ∼1mm craniotomy was performed either over the LGN (1.9mm lateral and 2.4 anterior from lambda). On the recording session, a multi-shank electrode (ASSY-37 E-1, 32-channels, Cambridge Neurotech Ltd., Cambridge, UK) was advanced to a depth of ∼3mm until visual responses to flashing stimuli were observed. Electrophysiology data was acquired with an OpenEphys acquisition board and units were isolated using Kilosort (Pachitariu et al., 2016a).

### Behavior and training

Mice ran through the corridor with no specific task (n = 4 animals, 65 sessions recording cortical activity; n = 3 animals, 19 sessions recording activity of LGN boutons). Prior to recording sessions, mice were placed in the virtual environment, typically for 3 days and for up to one week, until they were able to run for at least 80% of the time within a single session. 2 out of 7 mice ran without rewards. 5 out of 7 mice were motivated to run with rewards, by receiving ∼2.5 μl of water (4 mice) or of 10% sucrose (1 mouse) with the use of a solenoid valve (161T010; Neptune Research, USA). One animal received rewards at random positions along the corridor. The other 4 mice received rewards at the end of the corridor. For our experiments in cortex we chose various reward protocols to control for two things: 1. The effect of reward on our findings. Our results were the same regardless of whether animals received rewards or not. 2. The effect of stereotyped running speeds for mice receiving rewards at the end of the corridor. Our results were the same regardless of whether animals received rewards at a fixed position right before the end of the corridor or at varying positions along the corridor. Given the lack of influence by reward on our findings, rewards were placed at the end of the virtual corridor for subsequent recordings from LGN boutons.

For experiments in naïve animals (n = 2, Camk2a-tTA;tetO-G6s, Fig. 1e-h) mice were placed on the treadmill for 4 to 5 days, until they were able to run at speeds higher than 10 cm/s for at least 20 min. Only after animals reached this criterion we turned the VR on and started recording from the same field of view in V1 across multiple days.

We tracked the eye of the animal using an infrared camera (DMK 21BU04.H, Imaging Source) and custom software, as previously described (Saleem et al., 2018).

### Pre-processing of imaging data

Image registration in x and y, cell detection and spike deconvolution were performed with Suite2p (Pachitariu et al., 2016b). All subsequent analyses were performed on each neuron’s inferred activity from spike deconvolution, which we defined as the neuron’s ‘firing rate’. To account for the different decay times of the calcium indicators, the kernel’s decay timescale used for deconvolution was set to 0.5 s for GCaMP6f and to 2s for GCaMP6s.

For the LGN boutons data, we additionally used the method of Schröder et al. (2020) to align image frames in z-direction (along depth). By using a stack of closely spaced planes (1.8 µm inter-plane distance), we were able to detect small boutons across multiple planes, which could have otherwise moved outside a given plane due to brain movement in the z-direction. In brief, for each imaging stack, the algorithm estimates the optimal stack shift that maximizes the similarity of each plane to their corresponding target image (with target images across planes having been aligned to each other in x and y direction). After, assigning the shifted planes to their corresponding target image, a moving average across 2 to 3 neighboring planes is applied, resulting in a smooth image, and consequently in smooth calcium traces from boutons sampled from multiple, closely spaced planes.

Regions of interest (cell bodies or boutons) were detected from the aligned frames and were manually curated with the Suite2p graphical user interface, as described by Saleem et al. (2018). Data from V1 neurons with receptive fields in the periphery (>40°) are the same as in Saleem et al., (2018). These data were deconvolved and pulled together with data from V1 neurons with receptive fieds in the central visual field.

### Analysis of responses in Virtual Reality

To obtain response profiles as a function of position along the corridor, we first smoothed the deconvolved traces in time with a 250 ms Gaussian window and considered only time points with running speeds greater than 1 cm/s. We then discretized the position of the animal in 1 cm bins (100 bins in total) and estimated each neuron’s spike count and the occupancy map i.e. the total time spent in each bin. Both maps were smoothed in space with a fixed Gaussian window of 5 cm. Finally, each unit’s response profile was defined as the ratio of the smoothed spike count map over the smoothed occupancy map. We assessed the reliability of the response profiles based on a measure of variance explained and selected those with variance explained higher than 5%. To cross-validate the response profile patterns in Virtual Reality, we divided each session’s trials in two halves (odd vs even) and obtained a pair of response profiles for each unit. Odd trials were used as the train set, to determine the position at which cells or boutons preferred to fire maximally. Odd trials were subsequently excluded from further analysis.

The same splitting into odd and even trials was used to estimate each unit’s spatial modulation index (SMI). For each neuron or bouton, the position of the peak response was measured from the response profile averaged across odd trials (‘preferred position’). We then obtained the response, *R*_*p*._, at the preferred position and the visually identical position 40 cm away (‘non-preferred position’: *R*_*n*_), using the response profile averaged across even trials. Units with maximal response close to the start or end of the corridor (0-15 cm or 85-100 cm) were excluded, because their preferred position fell outside the visually matching segments. SMI was defined as:

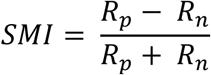

Therefore, a response with two equal peaks would have *SMI* = 0, whereas a response with one peak would have *SMI* = 1.

To cross-validate the response profile patterns and to estimate SMIs in passive viewing, we used the same odd trials from Virtual Reality as a train set. Based on those we obtained response profile patterns and SMIs from all trials during passive viewing. To isolate periods when the animal was stationary during passive viewing, we considered only time-points when the speed of the animal was less than 5 cm/s. Response profiles during stationary viewing were estimated only if the animal was stationary in at least 10 trials within a session. To isolate periods when the animal was running during passive viewing, we considered only time-points when the speed of the animal was higher than 1 cm/s. Response profiles in running during replay were estimated only if the animal was stationary in at least 10 trials within a session.

Reliability was defined as the variance in firing rate explained by the firing rate predicted by the cross-validated response profile. To predict firing rates, data were divided into 5 folds (5-fold cross-validation) and firing rates for each fold were predicted from responses profiles estimated from all other folds (training data). Reliability was defined as:

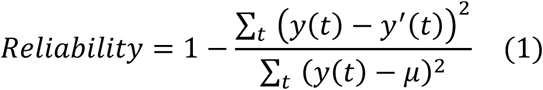

where *y*(*t*)is the actual, smoothed firing rate of the neuron at time *t where t is a time point between the beginning and end of the experiment, y*′(*t*)is the predicted firing rate for the same time bin, and *μ* is the mean firing rate of the training data.

The response reliability reported was obtained from the mean reliability across folds. Only neurons or boutons with response reliability greater than 5% were considered for further analysis.

### General linear models

To assess the joint contribution of all visual and behavioral factors in Virtual Reality we fitted the V1 responses to three multilinear regression models similar to Saleem et al. (2018). The models had the form: 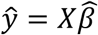,where *X* is an T-by-M matrix with T time points and M predictors and *ŷ* is the predicted calcium trace (T-by-1 array). Optimal coefficient estimates 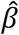 (M-by-1 array) that minimize the sum-squared error were obtained using: 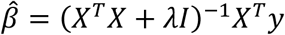, where *λ* is the ridge-regression coefficient.

The simplest model, the *visual* model, relied only on ‘trial onset’ (first 10 cm in the corridor), ‘trial offset’ (last 10 cm in the corridor) and the repetition of visual scenes within the visually matching segments (from 10 to 90 cm in the corridor). The basis functions for all predictors were square functions with width of 2 cm and height equal to unity. To model the repetition of visual scenes, a predictor within the visually matching segments comprised of two square functions placed 40 cm apart. Thus, the visual model’s design matrix had 30 predictors plus a constant: 5 predictors for trial onset, 5 predictors for trial offset and 20 predictors within the visually matching segments.

The second model, the *non-spatial* model was used to assess the influence of all the behavioral factors we measured: running speed, reward events, pupil size and the horizontal and vertical pupil position. These factors were added as predictors to the design matrix of the *visual* model, as follows: running speed was shifted backward and forward in time twice, in 500 ms steps, thus contributing 5 continuous predictors; pupil size and horizontal and vertical pupil position contributed 1 continuous predictor each; each reward event contributed one binary predictor at the time of the reward. The continuous predictors of running speed and pupil size were normalized between 0 and 1, whereas pupil position was normalized between −1 and 1 to account for movements in opposite directions.

The third model, the *spatial* model, allowed for an independent scaling of the two visually-matching segments in the corridor. For each predictor within the visually-matching segments, the two square functions were allowed to vary their height independently. The height of the two square functions was parameterized by a parameter *α*, such that the two functions had unit norm. An α = 0.5 corresponded to a purely visual representation with SMI close to 0, while *α* = 1 or *α* = 0 would correspond to a response only in the first or second segment, and an SMI close to 1.

To choose the best model, we used the ridge regression coefficient, *λ* that maximized the percentage of variance explained using five-fold cross-validation, searching the values *λ* = 0.01, 0.05, 0.1, 0.5 or 1. In the spatial model, we performed multiple ridge regression fits, searching for the optimal value of *α* using a step size of 0.1, for each *λ*.

The predictions obtained in the time domain were subsequently processed similarly as the original deconvolved traces, to obtain predicted response profiles and SMI.

